# 100 years domestication of penaeid shrimp and meta-analysis of breeding traits

**DOI:** 10.1101/2024.06.22.600213

**Authors:** Shengjie Ren, José M. Yáñez, Ricardo Perez-Enriquez, Morten Rye, Ross D. Houston, David A. Hurwood, Jose R. Gonzalez-Galaviz, Marcela Salazar, Dean R. Jerry

**Author notes:** Corresponding author: Shengjie Ren E-mail addresses.

## Abstract

Penaeid shrimp farming plays a pivotal role in ensuring future food security and promoting economic sustainability. Compared to the extensive long history of domestication observed in terrestrial agriculture species, the domestication and selective breeding of penaeids are relatively recent endeavors. Selective breeding aimed at improving production traits holds significant promise for enhancing efficiency and reducing the environmental impact of shrimp farming, thereby contributing to its long-term sustainability. Assessing genotype-by-environment (G-by-E) interactions is essential in breeding programs to ensure that improved penaeid shrimp strains perform consistently across different production environments, with genomic selection proving more effective than sib-testing alone in mitigating environmental sensitivity. Genome editing tools like CRISPR/Cas9 offer significant potential to accelerate genetic gains in penaeid shrimp by enabling rapid introduction of desired genetic changes, with recent advancements showing promising results in achieving high transfection efficiency in shrimp embryos. Additionally, artificial intelligence and machine learning are being leveraged to streamline phenotyping and enhance decision-making in shrimp breeding and farming, improving efficiency and accuracy in managing traits and predicting disease outbreaks. Herein, we provide an overview and update on the domestication of penaeid shrimp, including the current status of domestication for principal farmed species, key milestones in domestication history, targeted breeding traits in selective breeding programs, the advantages of integrating genomeic selection for enhancing production traits, and future directions for selective breeding of penaeid shrimp.

## Domestication of aquaculture species

### Comparison between agriculture and aquaculture

Historically, the process of domesticating aquaculture species differs remarkably from that of terrestrial agriculture. The majority of modern agricultural plants and animals were domesticated ca. 12,000 years ago, marking a pivotal moment in human civilization known as the Neolithic revolution^1-4^. In contrast, the domestication of aquaculture species is relatively recent^3,5^. It is estimated that around 543 aquaculture species have been domesticated since the early 20th century, with approximately 110 aquatic taxa domesticated since the 1980s^3,5,6^. Most of these species are still undergoing domestication, remaining close to their wild ancestors, often experiencing gene flow from wild populations, with domestication and selective breeding occurring concurrently. Certain aquaculture species, however, such as the common carp (*Cyprinus carpio*) in China and tilapia (*Oreochromis spp.*) in Egypt, show evidence of controlled reproduction dating back to around 1,500 years BC^7-9^.

While land agriculture sectors predominantly rely on a limited number of mammals, birds, and plant species, aquaculture production encompasses a remarkably diverse array of aquatic species. Currently, three key species—cattle, pigs, and chickens—contribute to 80% of global meat production, and four plant species—rice, maize, wheat, and potatoes—account for two-thirds of global plant production. In contrast, approximately 70 farmed aquatic species support 80% of worldwide aquaculture production^5,10^.

Moreover, the success rates of domestication vary significantly between aquatic and terrestrial species. Despite a much longer history spanning approximately 12,000 years, the domestication of land species has been considerably less successful. Only 0.08% of known land plants and 0.0002% of known animals have been successfully domesticated^11^. In contrast, domestication efforts for aquatic species have been notably more effective, with success rates of 0.13% for known aquatic animal species and 0.17% for known aquatic plant species^11^. Nevertheless, the rate of domestication for aquatic species is rapidly increasing, approximately 100 times faster than the rate observed for land species^3^.

### Levels of domestication for penaeids

In this manuscript, domestication is defined as the process involving the control of wild species for reproduction, the completion of part or full life cycles in captivity, and the utilization of modern genetic breeding techniques to enhance production traits. Accordingly, the classification of domestication is determined by the extent of human control over the life cycle of farmed species in captivity^6,9^. Typically, five levels of domestication ranging from 1 (least) to 5 (most), with an additional level 0 (involving only the exploitation of wild resources), are utilized for assessment^12,13^. Levels 1, 2, and 3 are classified as the pre-domestication stage, during which aquaculture activities still partly or fully rely on wild populations (See **Figure 1**.). In levels 4 and 5, the complete life cycle has been successfully managed from adult to egg to adult in captivity without the need for input from wild populations, indicating true domestication (See **Figure 1**.).

**Figure 1.**
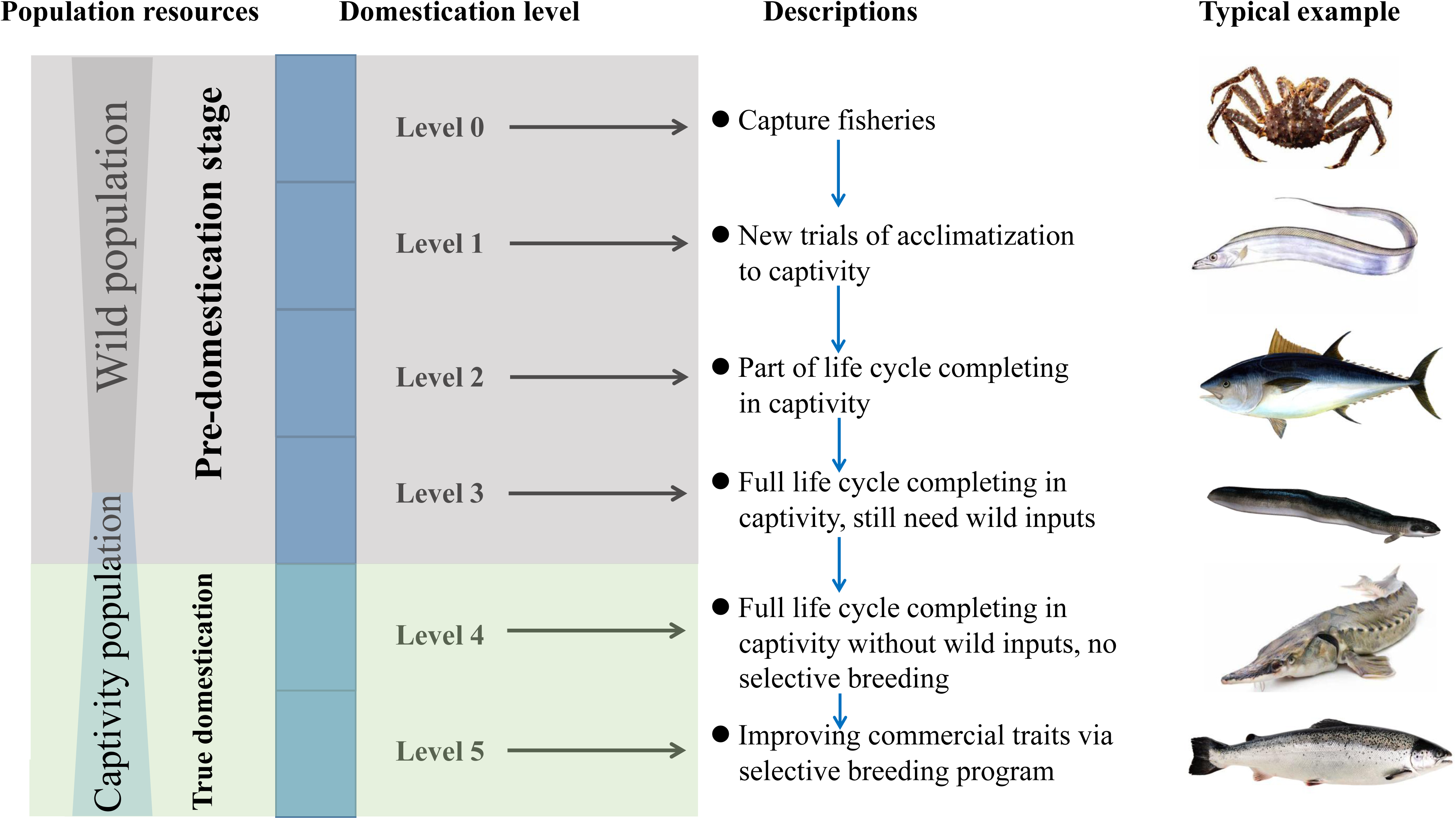
Domestication levels for aquaculture species.

Despite there being more than 110 shrimp species across 12 genera within the Family Penaeidae^14^, today’s farmed shrimp industry heavily relies on seven key species: *Litopenaeus vannamei, Penaeus monodon, Marsupenaeus japonicus, Fennerropenaeus chinensis, F. indicus, F. merguiensis,* and *L. stylirostris*. These principal penaeid species account for 98.2% of the global farmed shrimp production, which reached 7.24 million tons in 2021 (**Table 1**). Particularly noteworthy are *L. vannamei* and *P. monodon*, which play dominant roles in farmed penaeid production, constituting up to 97% of the global farmed shrimp production.

**Table 1.**
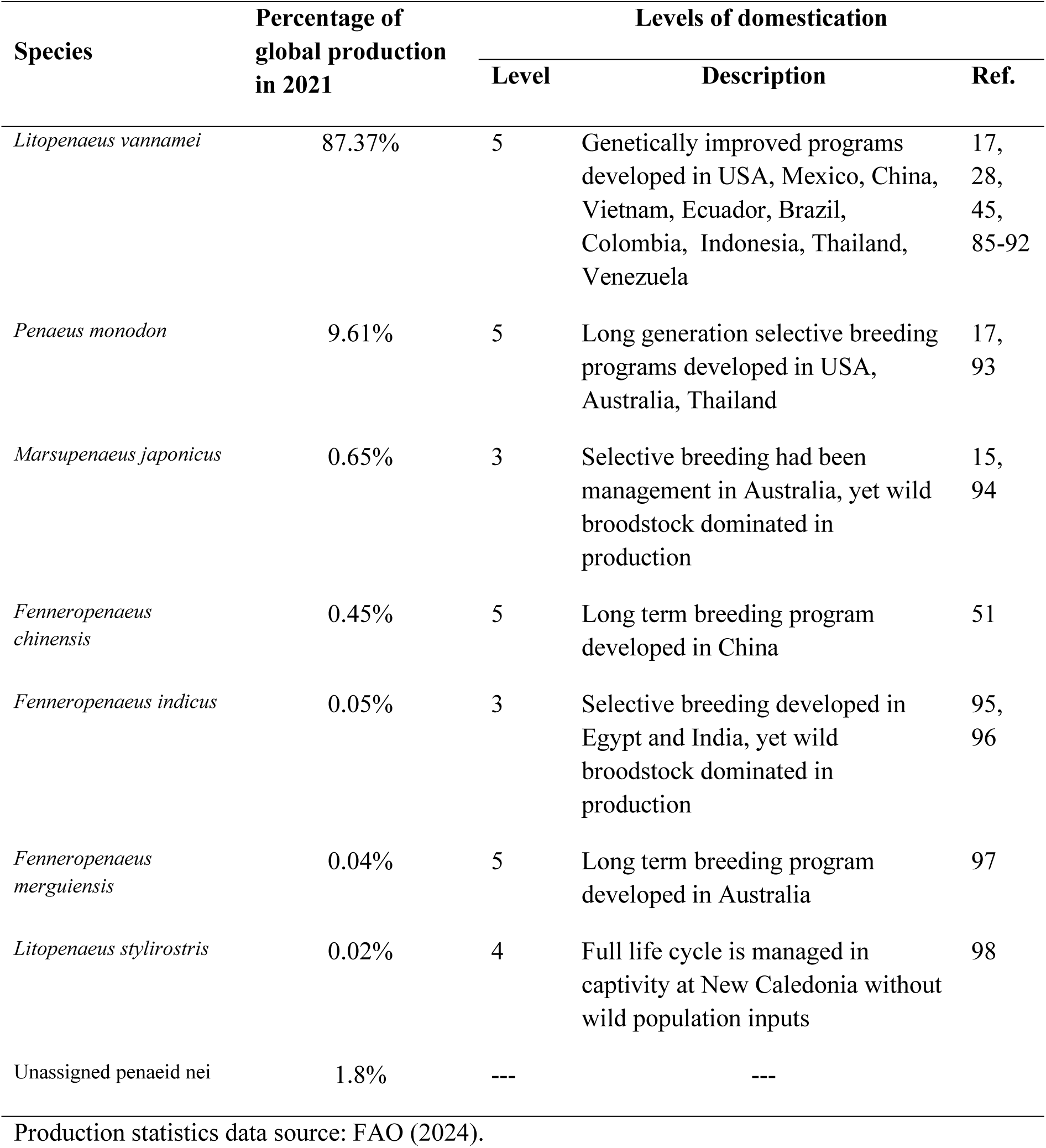
Domestication level of principal farmed Penaeid shrimp.

| Species | Percentage of global production in 2021 | Levels of domestication |  |  |
| --- | --- | --- | --- | --- |
|  |  | Level | Description | Ref. |
| *Litopenaeus vannamei* | 87.37% | 5 | Genetically improved programs developed in USA, Mexico, China, Vietnam, Ecuador, Brazil, Colombia, Indonesia, Thailand, Venezuela | 17, 28, 45, 85-92 |
| *Penaeus monodon* | 9.61% | 5 | Long generation selective breeding programs developed in USA, Australia, Thailand | 17, 93 |
| *Marsupenaeus japonicus* | 0.65% | 3 | Selective breeding had been management in Australia, yet wild broodstock dominated in production | 15, 94 |
| *Fenneropenaeus chinensis* | 0.45% | 5 | Long term breeding program developed in China | 51 |
| *Fenneropenaeus indicus* | 0.05% | 3 | Selective breeding developed in Egypt and India, yet wild broodstock dominated in production | 95, 96 |
| *Fenneropenaeus merguensis* | 0.04% | 5 | Long term breeding program developed in Australia | 97 |
| *Litopenaeus stylirostris* | 0.02% | 4 | Full life cycle is managed in captivity at New Caledonia without wild population inputs | 98 |
| Unassigned penaeid nei | 1.8% | --- | --- |  |

Production statistics data source: FAO (2024).

In the 1980s, the full life cycle of seven key farmed shrimp species was successfully closed in captivity, indicating domestication levels exceeding Level 3 (**Table 1**). However, most of these domesticated populations have not been maintained for long generations. In reality, *P. monodon* was not properly domesticated until nearly 2010 and to the point in the production of larger numbers of families that selective breeding programs were possible (D. Jerry pers. comm). Among penaeid species, *L. vannamei* stands out, by far, as the most advanced in domestication, boasting numerous selective breeding programs targeting various production traits. This progress is evident in the establishment of numerous pedigree selection-based programs globally (**Table 1**). Because domesticated strains offer a more reliable seed supply, many countries have initiated genetic improvement projects specifically for *L. vannamei*. Consequently, commercial seed supply for this species relies almost entirely on genetically enhanced broodstock. In the case of *P. monodon*, whilst the domestication level is now assessed as 5, supported by three selective breeding lines in the USA, Australia, and Thailand, the breeding of gravid female wild stocks of *P. monodon* remains prevalent in most Asian hatcheries. However, improvement of domestication levels in penaeid shrimp remains challenging. For instance, Australia established the first selective breeding program for *M. japonicus* targeting the lucrative Japanese market, but this program is currently defunct^15^. Apart from *L. vannamei,* the hatchery sector for seed production still heavily relies on gravid female wild stocks for the other six shrimp species. Furthermore, only one long-term selective breeding program for *F. chinensis* is underway in China, elevating the domestication level of this species to level 5.

Changes in production volume for key farmed penaeid species over time are significantly influenced by the domestication history of penaeid shrimp (**Figure 2**). Dr. Motosaku Fujinaga pioneered shrimp farming technology for *M. japonicus*, focusing on maturation and semi-intensive postlarvae (PL) production. During the 1950s to the late 1960s, *M. japonicus* played a dominant role, marking the inception of shrimp farming. The second wave of shrimp farming development occurred with the adaptation of *M. japonicus* maturation and PL production technology to *P. monodon* by Dr. I-Chiu Liao. This adaptation catalysed a significant expansion in shrimp farming across Asia from the late 1960s to the 2000s^16^.

**Figure 2.**
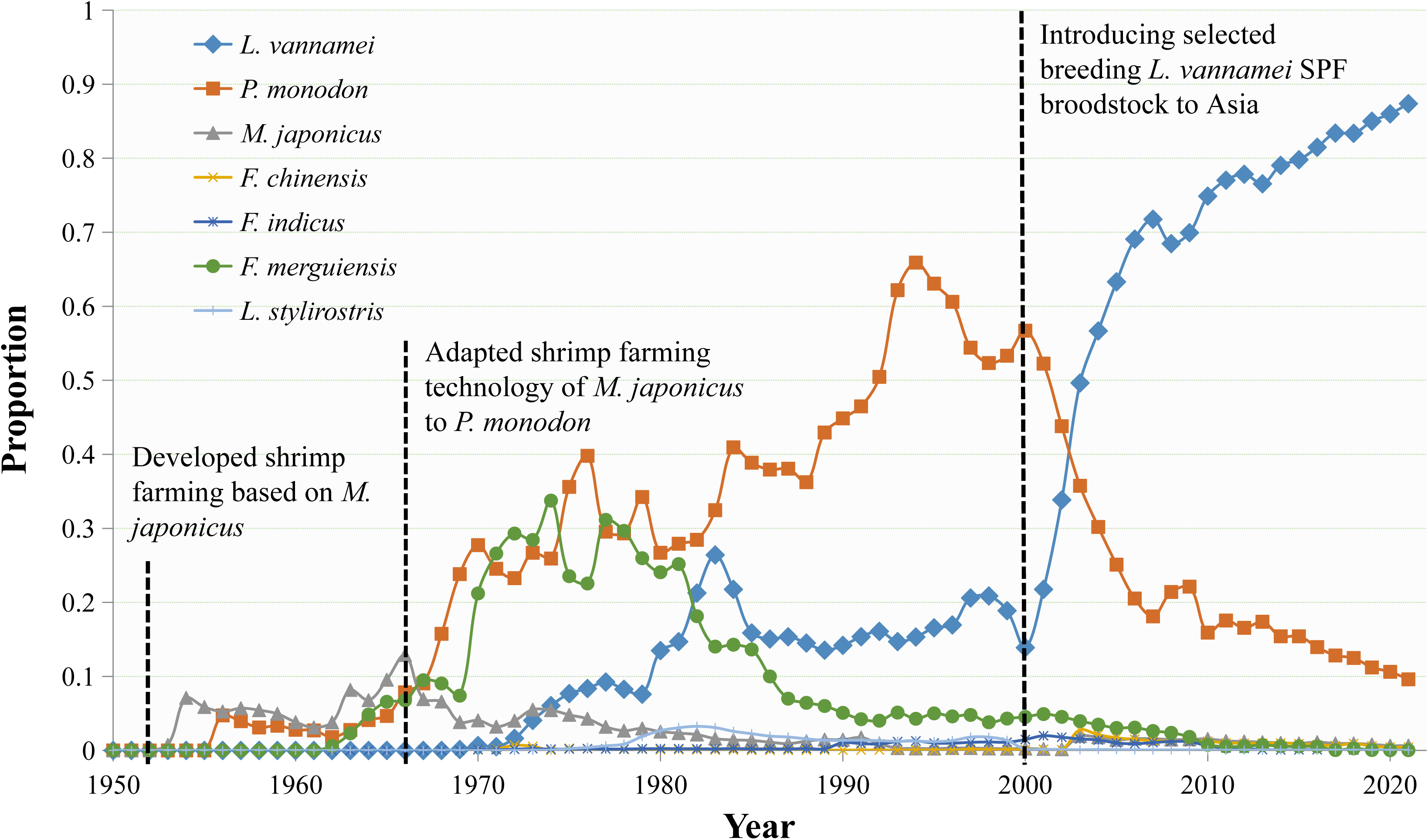
Trends of proportion of the production for principal farmed penaeid shrimp species in global shrimp farming production from 1950 to 2021.Production data statistics are from FAO 2024.

The development of hatchery and grow-out production of *F. merguiensis, L. vannamei,* and *F. chinensis* followed just a few years after the development of *P. monodon* as an aquaculture species. Consequently, these four species have played pivotal roles in farmed shrimp production during this period. However, data sources do not include production figures for *F. chinensis*, as China was the primary producer of this species, and its production statistics were often amalgamated with those of other penaeid species. More detailed information on production trends for *F. chinensis* during this time can be found in Benzie (2009)^17^. Since 2000, a remarkable growth in farmed shrimp production has been achieved by introducing genetically improved *L. vannamei* specific pathogen-free (SPF) broodstock to Asia (**Figure 2**). Within five years of introducing SPF *L. vannamei* to Asia, its production surpassed that of *P. monodon*, establishing *L. vannamei* as the dominant farmed penaeid species.

## The milestones of penaeid shrimp domestication

The earliest recorded trials of acclimatizing penaeid shrimp to farm conditions date back centuries in China, particularly in the practice known as ‘jiwei’ shrimp farming. The term ‘jiwei’ originates from Cantonese, used in Hong Kong, referring to ponds with dams in tidal zones. Farmers would introduce wild shrimp fry, along with milkfish, mullet, and other coastal finfish fry, into these ‘jiwei’ ponds via the pond dam during spring migrations into tidal impoundments^18^. Typically, ‘jiwei’ shrimp farms yield low production levels, ranging from 100 to 200 kg/ha/year of incidental crops, as there are minimal additional inputs until harvesting. It is noteworthy that in many coastal regions of China, ‘jiwei’ shrimp remains a common term for several penaeid shrimp species, including *Marsupenaeus japonica* and *M. ensis*. Similarly, relying on blue shrimp (*Litopenaeus stylirostris*) PL captured from the natural environment, the first shrimp farming in Mexico began in 1967 at Puerto Peñasco, Sonora, in the northwest of Mexico. This extensive type of shrimp farming remained unchanged until the 1980s. However, the development of penaeid shrimp domestication technology progressed slowly until the early twentieth century, primarily due to the complexity of their life cycles. Generally, the full life cycle of these species involves oceanic reproduction, a complex series of larval developmental stages, and an estuarine juvenile phase.

The first significant step towards closing the life cycle of penaeid shrimp in captivity occurred in 1934 when Dr. Motosaku Fujinaga^19^, based in the Yamaguchi Prefecture of Japan, successfully induced mature *M. japonicus* females to spawn in captivity and reared the nauplii to the subadults. A few years later, he succeeded in rearing *M. japonicus* larvae to adults^20^. However, his further domestication experiments were unfortunately suspended due to the outbreak of World War II. After the war, Dr. Fujinaga and his colleagues developed techniques for spawning gravid female shrimp, established feed protocols for larval rearing, and introduced semi-intensive grow-out technology, laying the groundwork for the modern shrimp farming industry. Even today, in many countries, the reliance on ready-to-spawn gravid female wild stocks remains widespread as a source of nauplii (i.e. particularly for *P. monodon*).

The second wave of penaeid shrimp domestication occurred in the 1960s, marked by the transfer of Fujinaga’s *M. japonicus* method to other penaeids and locations. One notable development took place at the Galveston Laboratory in the America, where several Gulf of Mexico penaeid species, including *P. aztecus, P. duorarum,* and *P. setiferus* had their life-cycles successfully closed through modifications of the Japanese culture methods^21^. Larval rearing method, known as the "Galveston Method" or "Clearwater Method," serves as the prototype for modern penaeid shrimp hatcheries. This method entails an intensive larval culture system, typically featuring indoor conical tanks, aeration airlifts, marine algae culture units, Artemia hatchery tanks for feeding mysis and postlarvae, and the reduction of water environment metal toxicity using EDTA. It ensures intensive and reliable production for larval culture. The Galveston Laboratory was also recognized as a pivotal research and training center for penaeid shrimp maturation-hatchery aquaculture experts worldwide from the 1960s to the early 1980s^22^.

The commercial shrimp farming industry took root in Taiwan when Dr. Liao transferred the spawning and larval rearing methods from *M. japonicus* to *P. monodon* after completing a postdoctoral fellowship at Fujinaga’s laboratory. Upon returning to Taiwan in 1968, Dr. Liao established the Tungkang Marine Laboratory and adapted Japanese penaeid domestication technology to six species: *P. monodon, L. stylirostris, P. penicillatus, M. japonicus, P. semisulcatus,* and *M. sp*^23,24^. After comparing the performance of these species on farms, *P. monodon* emerged as the most promising candidate for shrimp farming, leading to the rapid expansion of the industry. Under Dr. Liao’s direction at the Tungkang Marine Laboratory from 1971 to 1987, annual shrimp production skyrocketed from 427 tons to 88,264 tons, marking a staggering 200-fold increase.

A modification method of larval rearing, known as the "Taiwanese Method," played a crucial role in the development of modern shrimp farming in Asia. During the 1970s to 1980s, numerous Taiwanese technicians disseminated information about *P. monodon* farming across Southeast Asia. The first introduction of the "Taiwanese Method" to mainland China occurred into Fujian province, where local hatchery technicians adapted these technologies and developed a highly intensive larval rearing method known as the "Fujianese Method." Today, technicians from this region continue to employ the "Fujianese Method," which remains a significant role in the hatchery sector of shrimp farming in China.

Closing the full life cycle of penaeid shrimp in captivity remained challenging until the mid-1970s when maturation technology involving unilateral eyestalk ablation was first developed by a French research group in Tahiti^25-27^. Despite the first report on eyestalk ablation for manipulating hormones to induce ovarian maturation in crustacean species dates back to 1943^28^, it took three decades to apply this technique to penaeid shrimp. Early experiments demonstrated that eyestalk ablations could stimulate ovarian maturation, but female shrimp typically reabsorbed their ovaries or experienced lethal disruptions rather than spawning^26,29^. Subsequent research addressed these challenges using unilateral eyestalk ablation, which provided a moderate hormonal stimulus for successful spawning^30,31^. Due to improving animal welfare standards and associated certifications, currently, several leading shrimp breeding programs have recently developed broodstock production without eyestalk ablation in the most highly produced shrimp species. However, for fifty years now, unilateral eyestalk ablation has remained a standardized procedure for penaeid broodstock maturation and production of larvae worldwide. This not only bridged the final gap towards closing the life cycle of penaeid shrimp in captivity, but also paved the way for harnessing modern genetics in penaeid shrimp selective breeding programs.

The first genetic improvement project for Pacific white shrimp (*L. vannamei*) began in 1989 in Hawaii under the United States Marine Shrimp Farming Program (USMSFP)^32-36^. This selective breeding program initially adopted the concept of specific pathogen-free (SPF) from the swine and poultry industries in Europe for penaeid shrimp, aiming to produce high-health and genetically improved commercial broodstock or postlarvae (PLs) for cultivation^34,37^. The stocks were carefully curated through a stringent process involving the cautious screening of wild-caught shrimp. Individuals were selected based if they were naturally free from a predefined list of common shrimp pathogens. This selection process took place within a controlled quarantine environment at a nucleus breeding centre (NBC), housing numerous founding families. These selected stocks underwent a domestication and genetic improvement program, wherein the top-performing families from each generation were identified. PL from these families were then raised to become SPF broodstock in a highly secure broodstock multiplication center (BMC). Reflecting on the historical evolution of the shrimp farming industry, the development of genetically improved SPF breeding lines stands out as a significant technological advancement that shifted shrimp farming in Asia from *P. monodon* to *L. vannamei*, leading to the rapid expansion of shrimp farming globally^38^.

## Genetic improvement for penaeid shrimp

### Species and phenotype of interest

While selective breeding programs in agriculture began around 1900, drawing inspiration from Mendel’s pioneering hybridization experiments with peas in 1866^39^, it took another 15 years to formulate theoretical principles applicable to the genetic enhancement of farm animals^40^. However, modern breeding programs for aquatic species, particularly those employing large-scale family selection breeding designs, made their debut in aquaculture during the 1970s, notably in the context of salmonids^41^. The genetic improvement of penaeid shrimp has also seen significant attention, with 90 quantitative genetics papers published from 1997 to March 2024, highlighting the ongoing evolution and refinement of breeding strategies in penaeid species (**Table S1**). Within this body of literature, *L. vannamei* emerged as the most extensively studied species (62 publications), followed by *P. monodon* (11) and *F. chinensis* (7) (**see Figure 3**). Correspondingly, the leading contributors to these publications by nation were China (36), Mexico (21), and Australia (9). Notably, research from Mexico predominantly focuses on *L. vannamei*, while Australian studies concentrated on *P. monodon* and *F. merguiensis*. Conversely, Chinese research exhibits a broader interest, encompassing *L. vannamei, P. monodon, F. chinensis*, and *M. japonicus*. Among the eight studied species of penaeids, seven are tropical species, except for *F. chinensis*, which is a temperate species.

**Figure 3.**
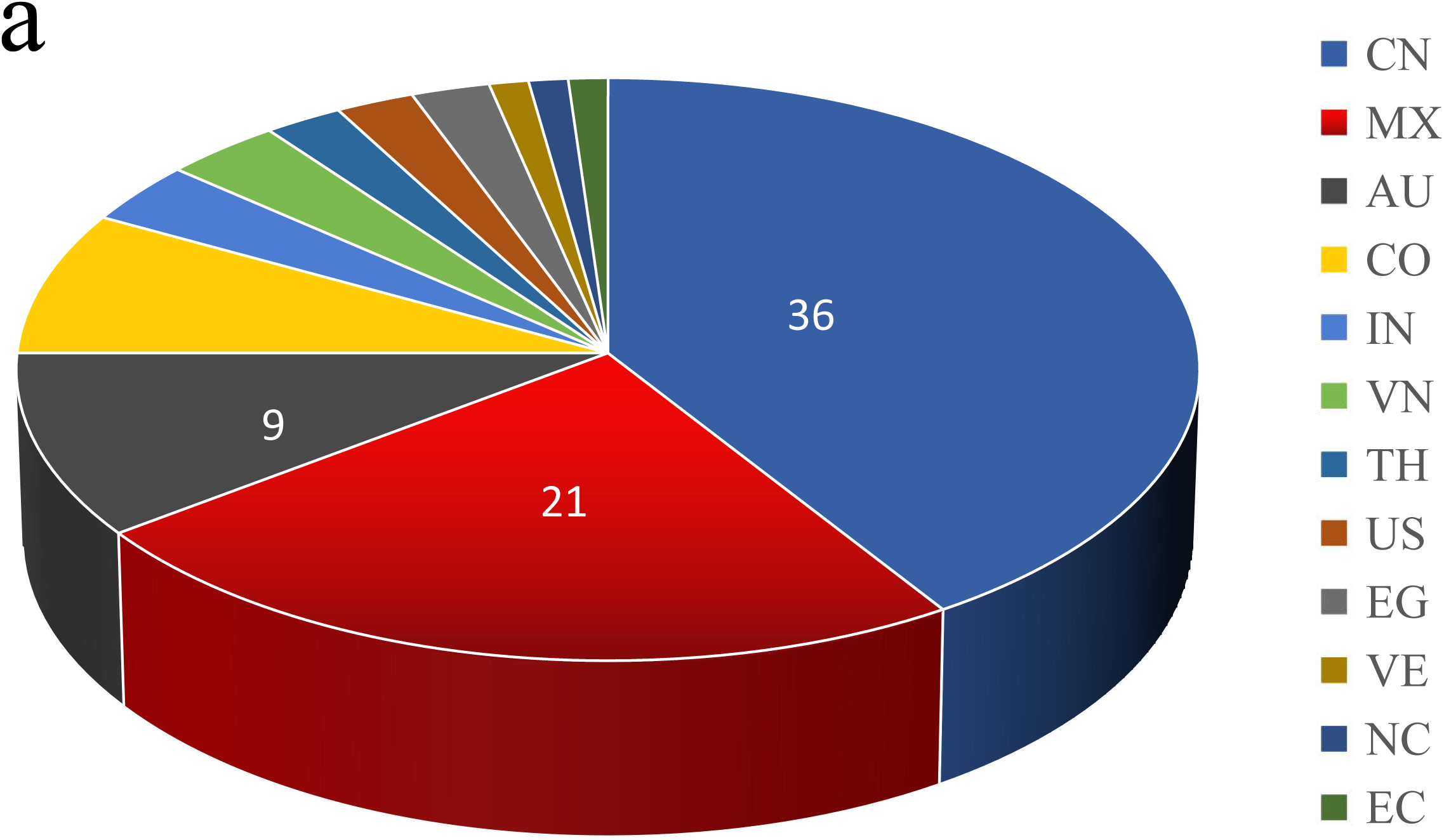
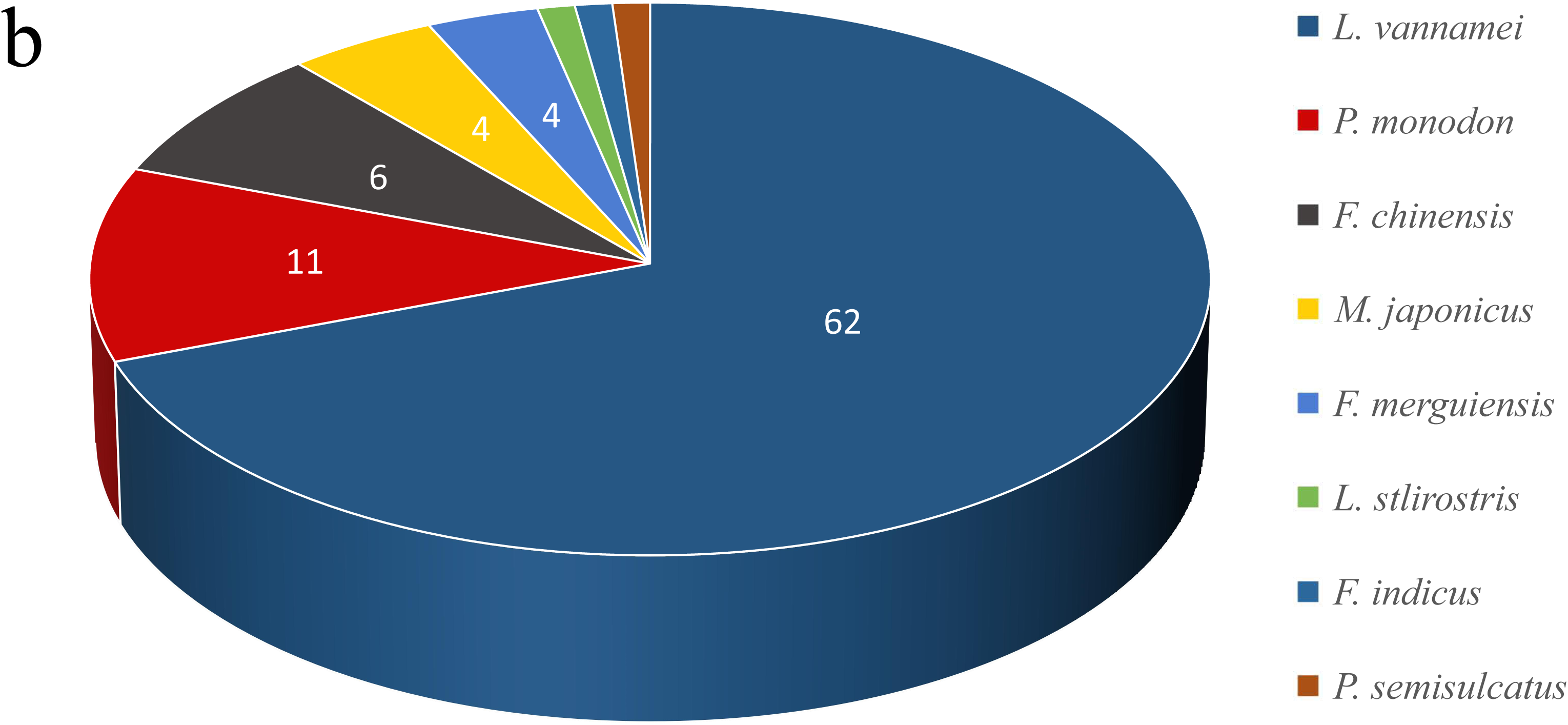
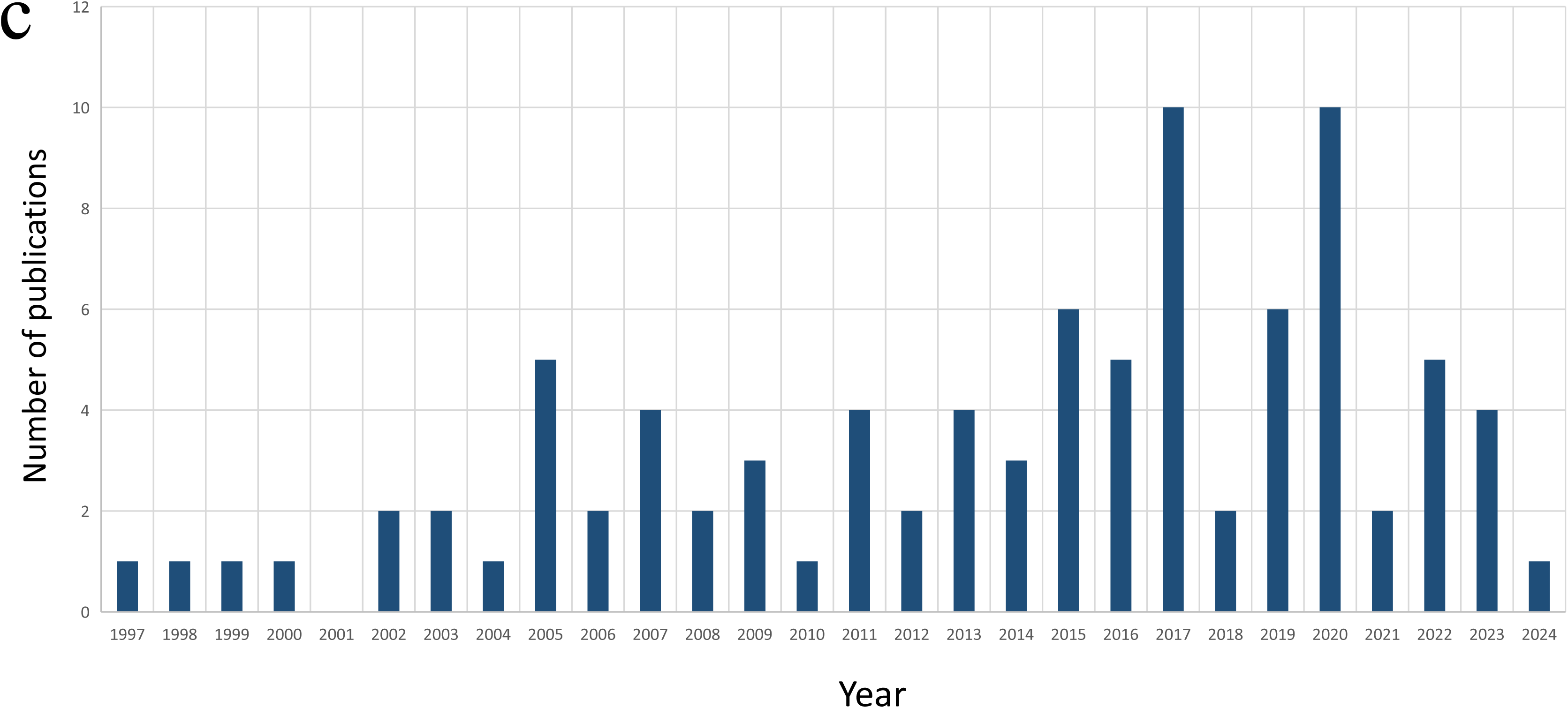
Overview of quantitative genetic publications for penaeids genetic improvement programs from 1997 to 2024. a, The leading contributors to these publications by nation. b, Number of quantitative genetic publications for per species of penaeid shrimp. c, Annual publications of quantitative genetic papers on penaeid shrimp selective breeding programs.

In genetic improvement programs, the breeding goals for phenotypic traits should align with the economic significance of culture traits that are heritable and accurately measurable^42,43^. Upon reviewing current genetic improvement programs for penaeid shrimp species, we identified 543 genetic estimates for various phenotypic traits (refer to **Table S2**). These traits of interest in selective breeding can be categorized into eight groups: growth traits, morphological traits, survival, disease resistance, stress tolerance, reproductive traits, quality traits, and feeding efficiency.

### Growth traits

From the farmers perspective, the growth rate stands out as one of the most economically significant traits. Enhancing this trait can result in more frequent harvests per year and/or larger individuals within the same cultivation period, thereby increasing market profits. Moreover, improvements in growth rate can indirectly positively impact other commercially relevant traits, such as feed conversion efficiency and survival rate^43-46^. Among 543 genetic estimates for the phenotypic traits of penaeid shrimp, growth-related traits are the most frequently investigated breeding traits, with 155 recorded estimates (see **Figure 4**).

**Figure 4.**
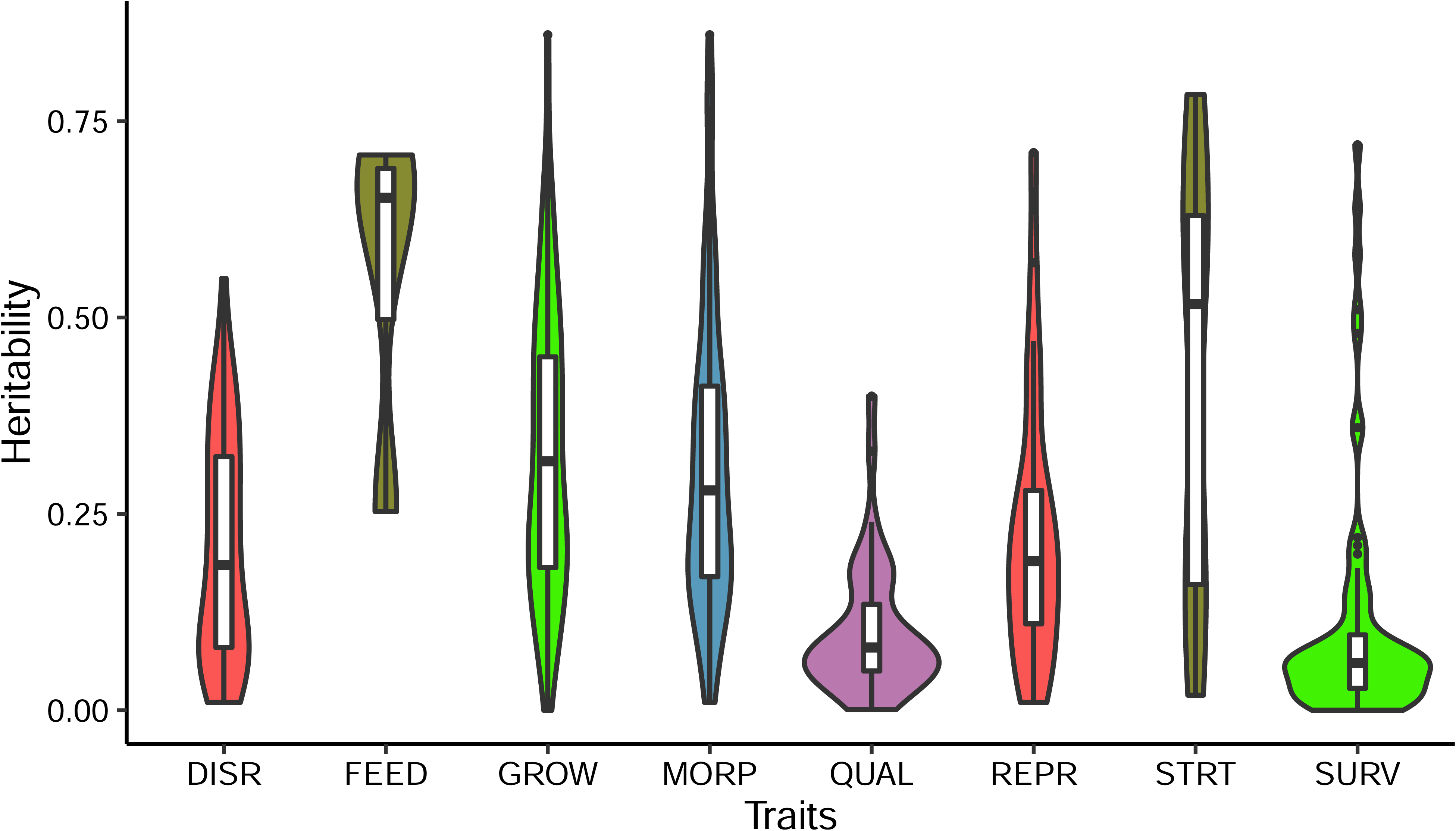
Analysis of heritability estimates on breeding traits in penaeids selective breeding programs. DISR, disease resistance; FEED, feeding efficiency; GROW, growth traits; MORP, morphological traits; QUAL, quality traits; REPR, reproductive traits; STRT, stress tolerance; SURV, survival.

While growth can be assessed through various definitions—such as absolute, relative, or specific terms—shrimp breeders commonly measure growth by monitoring ‘weight at age’, specifically focusing on body weight^47^. Our data analysis reveals that body weight (BW) is the predominant measurement for evaluating growth traits, whereas others include four instances using average daily gain (ADG) and three instances employing growth rate (GR). However, tail weight, net meat weight, meat yield, predicted net meat weight, and predicted meat yield are each used in only one case for assessing growth traits. A broad spectrum of heritability (*h²*) estimates for growth-related traits has been reported across penaeid species (see **Figure 4**). Among the 155 heritability estimates for these growth-related traits, the average *h²* was calculated to be 0.33 ± 0.02, ranging from 0.11 to 0.86 (see **Figure 4**). These results indicate substantial levels of additive genetic variation for growth-related traits, suggesting that artificial selection approaches will be effective for improving growth traits.

Due to the relatively high heritabilities observed in growth traits, coupled with the high fecundity and short generation intervals (1 year) of penaeids, numerous breeding programs have successfully attained notable genetic gains. For instance, Argue et al. (2002)^48^ reported that following a single generation of selection in *L. vannamei*, selected lines demonstrated a respective 21% and 23% accelerated growth compared to a control line devoid of selection, as evidenced in two distinct farming environments for raceways and ponds. Kenway et al. (2006)^49^ documented that the body weight of a selected line of *P. monodon* was 10% greater at 30 weeks of age, while Hetzel et al. (2000)^15^ reported genetic gains of 10.7% per generation for growth rate in *P. japonicus*. Additionally, Goyard et al. (2002)^50^ observed a selection response for *L. stylirostris* of 21% over five generations, while Sui et al. (2016)^51^ reported a selection response for *F. chinensis* of 18.6% over the same period. However, genetic improvement of growth rates must be carefully managed in practice. For instance, (i) common environmental effects (*c^2^*) due to the separate rearing of larvae from different crosses can be confounded with additive genetic effects, necessitating replication or communal rearing to address this issue, (ii) age must be properly controlled, and (iii) genotype-by-environment interactions should also be considered, as growth performance in one environment may only be moderately correlated with growth in another environment. Including a random common environmental effect (*c^2^*) into the statistical model for genetic evaluation can also be an alternative to estimate variance of the effect (*c^2^*) and correct it for a more accurate heritability estimation and estimate breeding value (EBV) prediction. Regarding age, it can be "controlled" or "measured" to correct body weight for it.

### Morphological traits

Morphological traits rank among the second most extensively studied breeding traits, with 152 estimates recorded. These traits exhibit a broad diversity, encompassing measurements of length, width, or height of various body parts. Notably, body length (BL), total length (TL), and abdominal length (AL) emerge as the top three traits evaluated within the morphological traits category (refer to **Table S2**). The overall heritability estimates for morphological traits closely parallel those of growth traits, with a mean value of 0.33 ± 0.02 (see **Figure 4**). While the levels of quantitative genetic variation for specific traits may vary among farmed populations and species, there are general consistencies observed^52^. High heritability for morphological traits lends support to Hill’s argument that "Heritabilities (*h²*) tend to be highest for conformational traits and mature size, typically 50 percent or more, and lowest for fitness-associated traits such as fertility"^52-55^.

### Survival traits

There are 71 estimates on survival traits, ranking third among the number of studied trait groups. In quantitative genetics, survival traits are considered fitness-associated, and heritability levels tend to be relatively low. This meta-analysis of survival trait estimates for penaeids selective breeding programs indicates an overall heritability of survival traits at a low value of 0.11 ± 0.02. Despite the crucial role survival rate plays in the success of shrimp farming^43,56^, the low levels of heritability observed for survival traits suggest that the response to selection for general survival traits is likely to be minimal. Consequently, improving pond survival rates via a family selection approach presents a significant challenge. Alternatively, selecting for disease resistance against the most serious diseases affecting penaeids serves as a complementary approach to enhancing overall survival rates in culture.

### Disease resistance traits

Diseases pose a significant challenge to shrimp production in aquaculture^57-59^, with some estimates of loss of 40% globally. The objective of selecting for disease resistance is to cultivate strains that inherently limit specific pathogen replication in the host, and therefore, show increased survival or lower pathogen burden in experimental and farming conditions^130^. This strategy is preferred by shrimp farmers as it eliminates the need for additional management efforts or investment in more sophisticated culture facilities, with the only additional cost being slightly higher prices for specific pathogen resistant (SPR) seed^57^. Compared to SPF, which is a health or biosecurity term indicating the absence of pathogens in seeds, SPR is a genetic term that refers to the selection for disease resistance^104^. This resistance can be specific to certain pathogens or their strains. However, some stocks may exhibit resistance to multiple pathogens while remaining susceptible to others. Furthermore, developing disease-resistant strains offers the advantage of minimal adverse impacts on the environment and public health compared to certain alternative measures, such as the use of antibiotics or chemical treatments^57^.

In total, there are 66 estimates regarding disease resistance traits, with white spot syndrome virus (WSSV) and Taura syndrome virus (TSV) resistance being among the most extensively studied. Some earlier studies reported that heritability (*h^2^*) estimates for WSSV resistance are lower and may involve a potential negative genetic correlation between growth and WSSV resistance^57,58,122^. Overall, the heritability estimates for disease resistance traits average at 0.21 ± 0.02, indicating moderate levels of additive genetic variation existing for disease resistance traits across most penaeid breeding programs. Consequently, this group of traits holds potential for effective improvement through a family selection approach. For instance, Argue et al. (2002)^48^ reported an 18.4% increase in survival from TSV infection for selected families of *L. vannamei* compared to a control line after just a single generation of selection. Over a three-year selection program, mean survival rates after TSV exposure rose by 24% to 37% in the selected line of *L. vannamei*^60^. After 15 generations of selection, researchers at the Oceanic Institute (Hawaii, USA) documented several families exhibiting 100% survival rates after TSV exposure^61^. Presently, TSV-resistant broodstock are extensively utilized in commercial hatcheries, rendering TSV no longer a significant threat to the global shrimp farming industry^36^. On the other hand, selection against WSSV has been challenging and is still one of the main pathogens in several countries.

The inability to directly measure disease resistance in selection candidates complicates the development of disease-resistant shrimp strains. To address this challenge, some hatcheries have adopted unconventional strategies, such as integrating survivors of disease outbreaks into the breeding nucleus^58,61^. Today, the use of genomic information in breeding programs through genomic selection enables the accurate prediction of disease resistance in selection candidates based on the phenotypes of their close relatives^5^, thereby enhancing disease resistance in aquaculture species.

Additionally, low-to-moderate negative or unfavorable genetic correlations between growth-related traits and disease resistance have been observed^36,48,87,122^. This complicates the simultaneous improvement of these traits within a breeding program, often necessitating the prioritization of one trait over the other. One potential solution is to develop and maintain separate lines for disease resistance and growth, and then produce crossbred animals for commercial purposes. However, this approach requires significant infrastructure and logistics, which can diminish the overall improvement achieved by each independent line.

### Quality traits

For quality traits, particular attention is directed towards color appearance, essential fatty acid composition, and the ratio of meat yield. In penaeids, the cooking color known as "redness" holds significant economic importance in the shrimp farming industry, as consumers generally prefer shrimp with a deeper red compared to lighter-colored ones. In Australia, the shrimp market utilizes a color scoring system to identify premium shrimp, with products exhibiting higher levels of "redness" often returning superior premium prices^62^. Among the 18 estimates concerning color appearance traits in penaeids, the overall heritability is calculated at 0.13 ± 0.03, ranging from 0.001 to 0.4. Despite the generally low heritability of this trait group, genetic variations in body color exist in some cases, making selection for this trait feasible.

Regarding fatty acids (12 estimates in total), the overall heritability is generally low at 0.08 ± 0.02, ranging from 0.03 to 0.18 (**Table S3**). However, certain fatty acids of interest, such as eicosapentaenoic acid (EPA) and docosahexaenoic acid (DHA), as well as the content of highly unsaturated fatty acids (HUFA), exhibit heritability values that suggest potential for genetic improvement^63^.

The economic value of shrimp is heavily influenced by meat yield, particularly tail weight^64,65^. Therefore, selecting for tail percentage in penaeids breeding projects could enhance profitability in the shrimp farming industry. Although there are six estimates regarding the percentage of tail weight, results suggest a relatively low level of heritability for this trait (**Table S2**). Consequently, incorporating tail percentage into selection criteria in penaeids breeding programs may not significantly enhance economic profitability.

### Reproductive traits

Penaeid shrimp species exhibit several distinctive reproductive characteristics. Upon maturation in hatcheries, a significant portion of females may spawn infrequently or not at all, while a smaller fraction of females spawn multiple times. Consequently, these prolific females likely contribute the majority of nauplii produced^66-69^. Traits such as high fecundity, increased spawning frequency for ablated females, a high ratio of mating success, rapid egg incubation rates, and enhanced survival rates from nauplii to the post-larval stage are highly desirable in the hatchery and nursery sectors. Among 25 heritability estimates of reproductive traits, a diverse array of traits within the reproductive trait group is evident. The most extensively studied traits include spawning frequency (SF), number of eggs per spawning (NE), number of nauplii per spawning (NN), and hatchery egg survival rate (HR).

On average, the heritability (*h^2^*) of reproductive traits is calculated to be 0.23 ± 0.04, indicating moderate levels of additive genetic variation for these traits (**Figure 4**). Specifically, a crucial set of female reproductive performance traits—including the number of eggs per spawn, number of nauplii per spawn, and multiple spawning capacity—are likely to benefit from genetic selection. However, limited additive genetic variation was observed for other reproductive performance traits, notably egg diameter, egg hatching rate, and the relative fecundity per weight (g) of individual broodstock females. Consequently, these traits are unlikely to be improved via genetic selection.

### Stress tolerance traits

Stress tolerance traits are group of traits directly associated to animal welfare in captivity. In animal breeding however, methods on measuring stress tolerance are challenging. For instance, the enzyme levels in blood samples, cortisol in plasma, lysozyme and immunoglobulin titres have been developed for stress indicators in selection response. The measurement accuracy of these stress indicators is expensive and time costly. In penaeids selective breeding, survival is a widely used stress indicator under the assumption that high responder families/groups would have higher survival rates when challenged with stressful environments. In total, there are 24 estimates of stress tolerance traits of penaeids; setting for stress tolerance environments include cold temperature, salinity tolerance, ammonia tolerance, and hypoxic tolerance^70-73^. It is noteworthy that heritability for stress tolerance traits is highest among the eight group of breeding traits of penaeids for 0.42 ± 0.06 (**Table S3**). Hence, stress tolerance traits could be effectively improved because of their high levels of additive genetic variation.

### Feeding efficiency traits

Feed costs constitute approximately 60% of production expenses in intensive shrimp culture. Thus, selecting for feed efficiency traits could enhance the profitability of the shrimp farming industry while also minimizing its environmental impact for sustainability. However, compared to poultry and the cattle industries, directly selecting for feeding efficiency in shrimp is challenging due to the difficulty of measuring individual feed consumption and efficiency in aquatic environments^74,75^. Consequently, feeding efficiency is the least estimated traits among the eight groups in selective breeding traits for penaeids, with only three traits investigated: residual feed intake (RFI), feeding efficiency ratio (FER), and daily feed intake (DFI)^76,77^. Heritability estimates of these traits are high, with a mean value of 0.56 ± 0.07, suggesting that while accurate measurements of feeding efficiency traits in penaeids are difficult, these traits, in most cases, exhibit high heritability and can be effectively improved by genetic selection.

## Application of genomic selection

The efficiency of family selection for penaeid shrimps can be significantly enhanced by employing SNP markers covering the entire genome to trace genomic relationships, as compared to traditional family selection based on pedigree information^78^. While genomic selection, utilizing molecular information, can fully exploit additive genetic variation for individuals within families, conventional pedigree family selection only captures 50% of the additive genetic variations for between-family variation. Established family selection breeding programs for penaeid shrimps, relying on pedigree management and routine measurement of phenotype traits, have successfully improved production for several species. Incorporating genomic selection into penaeid breeding programs offers substantial advantages, including maximizing genetic gains and minimizing inbreeding^79^. Zenger et al. (2019)^103^ reviewed the use of genomic selection in shrimp and highlighted the challenges and opportunities of this advanced selection approach.

A key component of applying genomic selection to penaeid selective breeding programs involves the development of genotyping platforms. Presently, single nucleotide polymorphisms (SNPs) Array platforms have been developed for *L. vannamei* (18k and 50k)^80,81^, and *P. monodon* (D. Jerry pers. comm). Several commercial SNP arrays are available for *L. vannamei*, including the Illumina Infinium 6k and the custom ThermoFisher Affymetrix Axiom 43k SNP array (Benchmark Genetics, Norway), also there are two SNP arrays for low and high density (Affymetrix Axiom 50k) to provide commercial services (The Center for Aquaculture Technologies, Canada). Additionally, genotyping by sequencing (GBS) techniques, including specific-locus amplified fragment sequencing (SLAF-seq), streamlined restriction site–associated DNA genotyping (2b-RAD), genotyping by target sequencing (GBTS), and diversity arrays technology sequencing (DArT-seq), have been successfully applied in *L. vannamei* and *F. merguiensis* improvement programs (**Table 2**).

**Table 2.**
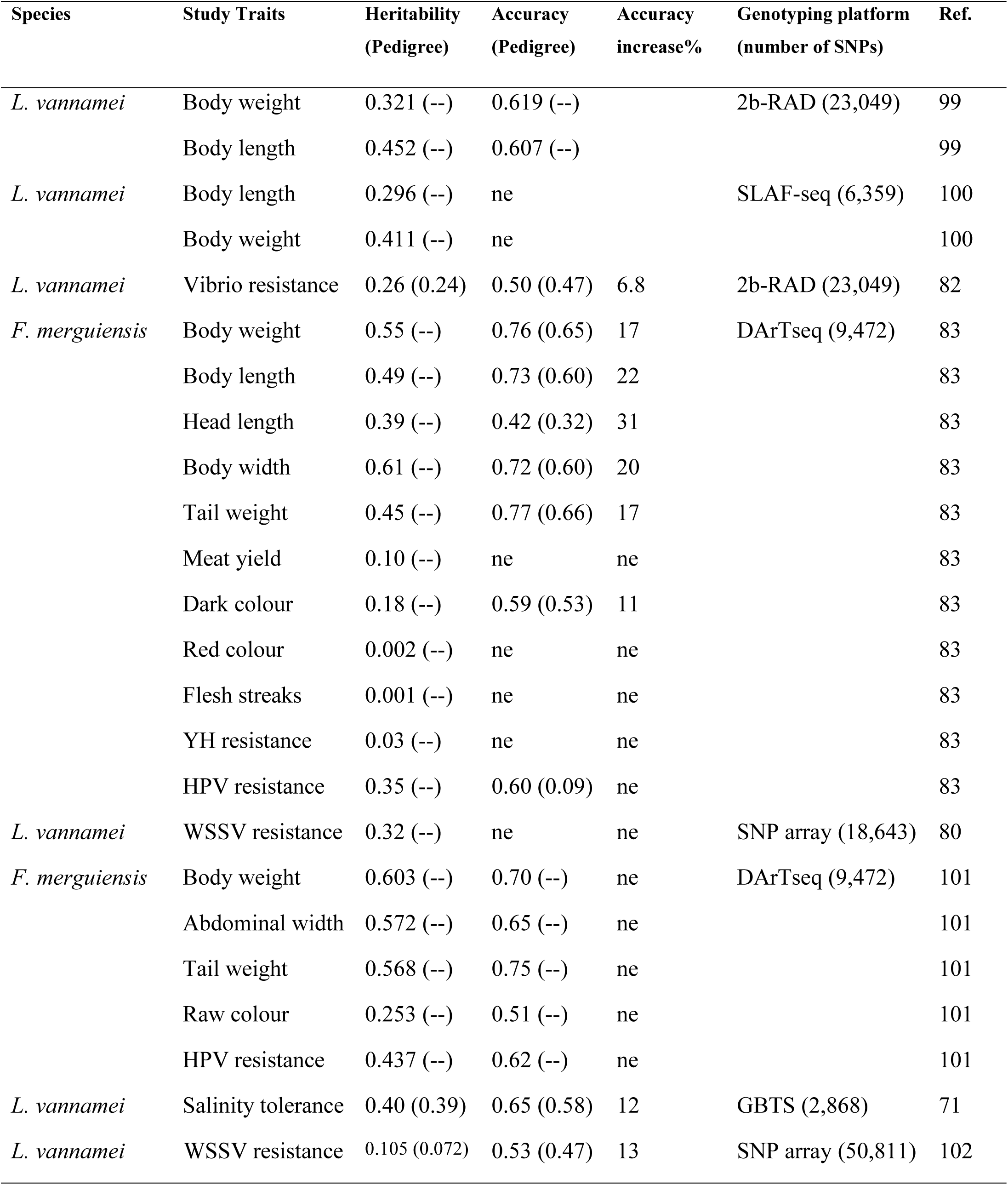
Summary on genomic selection study of Penaeid shrimp.

In summary, genomic selection for penaeid shrimps consistently demonstrates increased prediction accuracy compared to conventional pedigree selection across a range of traits. Studies report a median increase in prediction accuracy of 21.4% for growth traits, 9.9% for disease resistance, and 12% for salinity tolerance^71,82-84^. Moreover, these increases in prediction accuracy hold true across different species, genome evaluation models, and genotyping platforms, including both SNP arrays and GBS genotyping technology. The genomic best linear unbiased prediction (GBLUP) approach is the most widely used genomic selection model, utilizing genomic relationship matrices based on SNPs to estimate individual breeding values. While Bayesian models incorporating prior information of SNP effects have also been applied in penaeid breeding programs, prediction accuracies are generally comparable between GBLUP and Bayesian models.

Incorporating genomic selection tools into breeding programs for penaeid shrimp requires collecting genotype and phenotype information from thousands to tens of thousands of individuals in each breeding cycle, which can be particularly expensive. Advances in high-throughput genotyping technologies have improved the cost-effectiveness of applying genomic selection^105,106^. Additionally, the high costs of genotyping can be reduced through genotype imputation. This method involves genotyping a small number of parents with high-density (HD) SNP panels to serve as references while genotyping large numbers of progenies with low-density (LD) SNP markers, which can then be imputed to high-density SNP markers using the reference genotype information^107,108^. During the imputation process, statistical models, pedigrees, and a reference set of haplotypes (typically high-density genotypes from relatives) are utilized to infer missing genotypes for individuals genotyped using lower density chips. The application of genotype imputation in genomic selection has become increasingly popular, supported by various software packages such as Fimpute^109^, Beagle^110^, and AlphaImpute^111^. Since the accuracy of imputation is influenced by several factors, extensive research has been conducted to optimize analytical approaches and genotyping strategies. One key factor for accurate genotype imputation is the availability of a high-quality reference genome, which is the base to locate the molecular markers in their physical position. For instance, the current version of the reference genome for *L. vannamei* (ASM378908v1) is still at scaffold level (N50 = 605.6 kb), which difficult the accurate identification of the SNP coordinates into the chromosomes. Thus, genotype imputation is still difficult to implement in this and other shrimp species. However, recent advancements in dense linkage maps^136^ and future application of long-read sequencing techniques in *L. vannamei* will facilitate the implementation of cost-efficient genomic selection approaches by using genotype imputation from low-to high-density SNP panels.

## Genotype by environment interactions (G-by-E)

Ideally, improved seed developed in breeding programs should be more productive across a variety of commercial culture environments^112^. However, the relative performance of a specific animal phenotype depends on its genotype, the production environment, and the interaction between these factors. G-by-E interactions, or genotype by environment interactions, occur when the same genotypes exhibit different phenotypic responses under varying environmental conditions^112-115^. Therefore, assessing G-by-E interactions is crucial to determine whether an improved animal will perform consistently well in different production environments.

Studies on G-by-E interactions in penaeid shrimp species have focused on correlations between specific growth traits and various culture environments, particularly the effects of stocking density, location, and temperature. The general trend reported in the literature for penaeid shrimp is that genetic correlations for growth traits are very high, often close to 1.0, between similar environments, such as ponds, different farms within the same region, varying culturing densities, or growth performance at different ages^90,116-118^. However, when testing environments differ significantly, genetic correlations for growth performance tend to be low. For instance, correlations are low between low and optimal farming temperatures^119^, or between environments with and without a natural WSSV outbreak^120^.

To minimize the potential impacts of genotype-by-environment (G-by-E) interactions in practical selective breeding programs for penaeid shrimp, especially for international broodstock markets, the selection environment should closely mimic actual production conditions. Consistent measurements across different environments are crucial. Alternatively, selection should consider performance in both environments. Genomic selection can enhance the breeding of more robust strains by testing reference populations (full sibs) in diverse environments^121^. Evaluating genotype performance across an environmental gradient can help calculate G-by-E effects using genomic selection, which reduces sensitivity to environmental variation and yields significantly better results than sib-testing alone.

## Future directions

### Genome editing tools to accelerate genetic gains

Traditional selective breeding programs of penaeid shrimp rely on pedigree family selection or genomic selection approaches, which exploit the additive genetic variations present in the natural breeding population. Consequently, the potential genetic gains from selective breeding are constrained by the presence/not and the levels of heritability for the target traits within the nucleus population. In contrast, genome editing tools such as CRISPR/Cas9 can rapidly introduce desired changes to the genome, creating *de novo* alleles or incorporating alleles from other strains or species^123^. Recently, CRISPR/Cas9 genome editing has been successfully applied in vivo and in cell lines of several major aquaculture species, targeting traits such as sterility, growth, and disease resistance^5,123^. The high fecundity and external fertilization (artificial insemination) of penaeid species^68,86^ make them particularly suitable for genome editing research and applications on a scale that is not feasible in farmed terrestrial animals.

A key step for applying genome editing in penaeid shrimp is developing efficient methods for introducing the CRISPR/Cas9 system into cells or embryos. Traditional delivery strategies for CRISPR/Cas9 components have proven ineffective for genome editing in *L. vannamei* embryos. However, a new strategy using PEI-coated SWNTs nanocarriers has been developed to efficiently deliver CRISPR/Cas9 plasmids into early embryos of this species, achieving a transfection efficiency of 36%^124^. This study highlights an innovative approach for large-scale genome editing applications aimed at enhancing growth performance and disease resistance in penaeid shrimp breeding programs. Additionally, research has investigated three different cargoes—DNA plasmid, mRNA, and a recombinant protein of Cas9 system—for CRISPR/Cas9-induced gene editing in *L. vannamei* zygotes using both physical and chemical transfection methods^125^.

Innovative technology is crucial for advancing food production to meet the growing global demand. CRISPR/Cas9 technology holds exciting potential to enhance the quantity, quality, and sustainability of seafood production worldwide. However, its successful implementation depends on public and regulatory acceptance. There is significant debate regarding the definition of genetic modification (GM) and whether genome-editing approaches like CRISPR/Cas9 should be classified separately.

### Artificial intelligence (AI) and machine learning

One of the challenges for shrimp breeding programs is the requirement to phenotype for multiple traits thousands to 10’s thousands of individuals to accurately capture the variance in traits and to estimate genetic parameters and breeding values^103^. Historically this has been achieved by manual measurement, which is laborious, often invasive, and time-consuming. In addition, production of farmed shrimp is complex with farmers having to account for changing environmental parameters, disease, management processes and shrimp growth and biology. For both breeding programs and general production, the use of artificial intelligence based on computer vision, machine learning, and prediction are being developed to acquire phenotypic data more expediently and in the development of decision support software for farmers. For example, Saleh et al. (2024)^126^ used digital capture of 8,164 images and deep-learning training to predict 12 shrimp body landmarks in a breeding program for *P. monodon*. Prediction of these landmarks then was used to automate morphological and weight measurements. Similarly, Setiawan et al. (2022)^127^ used underwater cameras to capture images of *L. vannamei* and KNN regression machine learning to estimate the weight of live shrimp. In relation to calculation of genetic merit of shrimp in genomic-based breeding programs, machine learning has also been evaluated against different genomic selection models and shown to have potential to improve accuracy of prediction over GBLUP approaches^65^.

Machine learning approaches have also begun to be applied to unpack the complexity of shrimp farming and predict future events like disease outbreaks. For example, Khiem et al. (2022)^128^ and Tuyen et al. (2024)^129^ used various machine-learning models to predict the outbreak of diseases such as WSSV based on pond parameter datastreams. These models showed that training algorithms that incorporate environmental data has potential for farmers to be able to predict likelihood of disease events, offering them the potential to take management decisions earlier to limit impact.

### Genome-wide association studies for marker-assisted selection (MAS)

Genome-wide association studies (GWAS) have significantly advanced aquaculture breeding and genetics by identifying genetic markers associated with key traits, thus facilitating a better understanding on genetic variants controlling desirable characteristics and their implementation in breeding programs. In Pacific white shrimp (*L. vannamei*), GWAS has been instrumental in uncovering SNPs linked to various economically important traits, including resistance to white spot syndrome virus (WSSV)^102^, growth^131-133^, ammonia nitrogen tolerance^134^ and sex-determining region^135,136^.

For instance, a study involving a WSSV-resistant line of *L. vannamei* identified two SNPs significantly associated with survival post-infection, which explained a low genetic variance for the trait, suggesting a polygenic nature of WSSV resistance and highlighting the potential of genomic selection for this trait^102^. Additionally, GWAS on body weight (BW) and growth traits has pinpointed SNPs within or near candidate genes such as deoxycytidylate deaminase, non-receptor protein tyrosine kinase, and class C scavenger receptor (LvSRC), protein kinase C delta type and ras-related protein Rap-2a, which are associated with significant phenotypic variance in growth-related traits^131-133^. It has also been demonstrated that some of these genetic markers have enhanced the accuracy of marker-assisted selection (MAS) over traditional methods. Further, GWAS has identified critical genomic regions related to sex determination in *L. vannamei*^135,136^, *P. monodon*^137,138^, which is crucial for exploiting sexual dimorphism in shrimp growth rates, thus optimizing production efficiency. The identification of these markers and the understanding of their associated genetic mechanisms offer promising pathways for advancing selective breeding strategies, ensuring improved disease resistance, enhanced growth rates, and better tolerance to environmental stressors in shrimp aquaculture.

## Concluding remarks

Penaeid shrimp farming, plays a crucial role in ensuring food security, fostering economic sustainability, and addressing ecosystem service issues on a global scale. However, compared to the long history of domestication seen in terrestrial agriculture species, the process of domesticating and selectively breeding penaeids is relatively young. A significant milestone towards achieving closed life cycles of penaeid shrimp in captivity was reached in 1934 when Dr. Fujinaga successfully induced mature *M. japonicus* females to spawn. Today, the farmed shrimp industry heavily relies on seven key species, which collectively account for 98.2% of global farmed shrimp production. Successful closed full life cycle of these seven key farmed shrimp species in captivity was achieved in the 1980s.

Harnessing modern genetics in selective breeding programs for Specific Pathogen-Free (SPF) *L. vannamei* represents a remarkable technological leap that has revolutionized shrimp farming, facilitating its rapid global expansion. Genetic enhancement of production traits in penaeids primarily targets growth rates, morphological traits, survival rates, disease resistance, stress tolerance, reproductive capabilities, quality attributes, and feeding efficiency. Meta-analyses underscore the significant additive genetic variations observed in production traits, indicating promising results and substantial potential for selective breeding in penaeids, particularly concerning growth rates and disease resistance. The inclusion of genomic selection in penaeid breeding initiatives presents notable advantages, such as maximizing genetic improvements while minimizing inbreeding. Looking ahead, the commercial utilization of genome editing holds immense promise for enhancing economically significant traits in penaeid shrimp, thereby accelerating genetic gain and aiding in overcoming challenges faced by the shrimp farming industry.

## Data availability

All data used in this study is freely available via Zenodo at *************.

## Authors’ contributions

S.J.R. initially designed the study with substantial input from D.R.J., J.M.Y., J.R.G., M.R., R.D.H, M.S, R.P.E, and D.A.H. S.J.R. collected data and produced figures. All authors reviewed and provided inputs for the final manuscript. S.J.R. wrote the first draft manuscript with all authors subsequently providing input.

## Acknowledgements

S.J.R. acknowledges a fellowship position support by Biobreeding Insititute, Xianghu Laboratory.

## Additional files

### Additional file 1 Table S1

Format: csv file

Title: Quantitative genetic publications for penaeids genetic improvement programs from 1997 to 2024.

### Additional file 2 Table S2

Format: csv file

Title: Summary of heritability estimates on breeding traits in penaeids selective breeding programs.

### Additional file 3 Table S3

Format: csv file

Title: Analysis of heritability estimates in penaeids selective breeding programs for disaese resistance, feeding efficiency, growth traits, morphological traits, quality traits, reproductive traits, stress tolerance and survival.

